# Orthogonally targeted tumor radiosensitization using cell penetrating peptide-ATM inhibitor conjugates to stimulate anti-tumor immune responses

**DOI:** 10.64898/2026.01.09.698736

**Authors:** Kanika Dhawan, Michael M. Allevato, Jacqueline Lesperance, Maria F. Camargo, Marcus M. Cheng, Mahsa Mortaja, Bamdad Zareh, Dina V. Hingorani, Stephen R. Adams, J. Silvio Gutkind, Sunil J. Advani

## Abstract

Tumor resistance to radiotherapy continues to be a significant problem in improving cancer patient outcomes. To overcome radioresistance, drugs that sensitize cancer cells to ionizing radiation have been tested. In theory, radiosensitizers should increase irradiated tumor kill and improve patient outcomes. In practice, the clinical utility of such drugs is curtailed by radiosensitization of peri-tumoral normal tissues causing toxicities. To address these issues, we developed an activatable cell penetrating peptide-drug conjugate to deliver a small molecule radiosensitizer with spatial precision to tumors. The activatable cell penetrating peptide (ACPP) scaffold cloaks a cell penetrating peptide-drug conjugate until it is unmasked within tumors through matrix metalloproteinase cleavage. Using antibody-drug conjugate linker chemistry, we attached the potent ataxia-telangiectasia mutated (ATM) kinase inhibitor AZD0156 to ACPP and created ACPP-AZD0156. In immune-competent murine cancer models, tumor-targeted ACPP-AZD0156 in combination with ionizing radiation stimulated tumor immune infiltration by CD8+ T cells and increased tumor control when compared to non-targeted ATM inhibitor. Mechanistically, ACPP-AZD0156 radiosensitized tumor control was dependent on the adaptive arm of the immune system. Finally, the combination of radiotherapy and ACPP-AZD0156 potentiated immune checkpoint inhibitors that resulted in durable tumor control. The therapeutic synergies of ACPP targeted ATM inhibitor to radiosensitize and stimulate anti-tumor immune responses provides a rationale for developing tumor-targeted radiosensitizer drug conjugates that restrict radiosensitization to cancer cells that then engages anti-tumor immune responses to improve cancer patient outcomes.

## 1. Introduction

Over 50% of cancer patients receive radiotherapy as part of their treatment; however, intrinsic and acquired radioresistance limits its efficacy[1, 2]. To improve tumor control for patients with locally advanced cancers, radiotherapy is combined with concurrent cytotoxic chemotherapies such as cisplatin, taxanes, or 5-fluorouracil, which amplifies DNA damage and disrupt cell-cycle checkpoints [3]. Randomized clinical trials have demonstrated improved patient outcomes when combining chemotherapies with radiotherapy across diverse tumor types including glioblastoma, head and neck, esophageal, lung, bladder, rectal, anal and cervical cancers[4-11]. However, cytotoxic chemotherapies are dose-limited secondary to their systemic toxicities that constrains their ability to widen the therapeutic index of radiotherapy[12-14].

Cancer cell kill by radiotherapy is driven by ionizing radiation (IR) causing DNA double-strand breaks (DSBs) which is a lethal genomic injury that subsequently results in mitotic catastrophe and cell death[15, 16]. Mechanistic elucidation of the DNA damage response (DDR) and repair processes have identified druggable pathways that can achieve selective radiosensitization in irradiated cells compared to non-irradiated tissues[1, 17]. Among these, ataxia-telangiectasia mutated (ATM) kinase functions as a master regulator to coordinate DNA DSB repair. ATM phosphorylates histone H2AX (γH2AX) and downstream effectors including Chk2, BRCA1, and 53BP1 to mediate cellular responses to IR induced DNA damage[18-20]. Mutations in ATM result in exquisite radiosensitivity that can be pharmacologically recapitulated with ATM inhibitors (ATMi)[21-23]. A theoretical advantage of DDR inhibitor based radiosensitization compared to cytotoxic chemotherapies is that they are inherently less cytotoxic and selectively increase DNA damage in irradiated cells compared to non-irradiated cells. This provides a potential therapeutic window where systemically delivered DDR inhibitors increase DNA damage only in the irradiated target tissue but not in unirradiated normal tissues. However, their clinical translation of DDR inhibitors has been disappointing[13, 24]. Recent clinical trials combining DDR inhibitors with concurrent chemo-radiotherapy have shown they cause increased toxicity without improvement in tumor control[25-28]. One reason for the lack of clinical benefit of DDR inhibitors in irradiated cancer patients, is these free small-molecule inhibitors radiosensitize not only cancer cells within the intended tumor target but also peri-tumoral normal tissues that are located within the irradiated field. Thus, the benefit of radiosensitizing cancer cells is negated by collateral radiosensitized damage of adjacent normal tissues which may explain in part the clinical failures of DDR inhibitors to widen the therapeutic index of radiotherapy.

Within the last decade, immunotherapies have revolutionized cancer therapy with the understanding that immune checkpoints expressed on T-cells can result in tumors evading the immune system[29, 30]. Therapeutic targeting of immune checkpoints such as cytotoxic T-lymphocyte associated protein 4 (CTLA-4) or programmed death protein 1 (PD-1) overcomes tumor immunosuppression by reinvigorating exhausted T cell and has resulted in durable tumor control in a subset of cancer patients[31]. However, the majority of cancer patients are unresponsive to immune checkpoint inhibitors. To improve and broaden the efficacy of immunotherapies, they are being tested in combination with standard of care anti-cancer agents including radiotherapy[32-35]. Radiotherapy can complement immunotherapy by inducing immunogenic cell death and releasing tumor-associated antigens and damage-associated molecular patterns (DAMPs), activating the cGAS-STING pathway and type I interferon (IFN-1) signaling which enhance antigen presentation and prime cytotoxic CD8+ T-cell responses[36, 37]. Moreover, preclinical studies have shown that ATM inhibition in combination with IR increases tumor immune infiltration in syngeneic immune-competent cancer models [38-41].

To improve the therapeutic index of radiotherapy and immunotherapy, we hypothesized tumor-targeted ATM radiosensitizated immunogenic cell death could potentiate immune checkpoint inhibitors and achieve durable tumor control. As a solution to the non-selective distribution of systemically injected DDR inhibitors, we have developed an orthogonal spatially targeted strategy for tumor radiosensitization that combines the intrinsic advantage of DDR inhibitors to increase DNA damage in irradiated cells with spatially targeted tumor drug delivery. Here, we synthesized and performed preclinical investigations of a novel tumor-targeted activatable cell penetrating peptide-ATM inhibitor conjugate (ACPP-ATMi). This strategy combines the advantages of DDR pathway-specific radiosensitizers together with a tumor-targeted delivery vehicle. In systemic circulation, intact ACPP-ATMi cloaks the conjugated ATMi by making it inaccessible to cells. Within tumors, circulating ACPP-ATMi is unmasked by extracellular matrix metalloproteinases (MMP-2/9) resulting in tumor targeted uptake of the cell penetrating peptide-ATMi conjugate. Using preclinical syngeneic immune-competent cancer models, we show that tumor-targeted ACPP-ATMi increased IR-induced DNA damage, stimulated CD8+ T-cell infiltration within irradiated tumors, and improved tumor control compared to non-targeted ATMi. Moreover, ACPP-ATMi radiosensitization combined with PD-1 blockade resulted in durable tumor control. Taken together, our cloaked cell-penetrating peptide drug delivery platform supports a novel framework for precision DDR inhibitor radiosensitization for cancers by spatially targeting radiosensitizer delivery to stimulate anti-tumor immune responses that can then potentiates immunotherapies.

## 2. Material and methods

### 2.1 Cells and Reagents

Human A549 (CRM-CCL-185), LN229 (CRL-2611), CAL27 (CRL-2095), and murine LL2 (CRL-1642), B16 (CRL-6475), MEF (C57-BL/6-1, SCRC-1008) cell lines were obtained from American Type Culture Collection. Murine MC38 (ENH204-FP) cell line was obtained from Kerafast. A549, LN229, CAL27, LL2, and B16 cells were cultured in DMEM (Gibco) supplemented with 10% FBS (Omega Scientific). MC38 cells were cultured in DMEM supplemented with 10% FBS, 1mM sodium pyruvate (Gibco), 1% non-essential amino acids (Gibco), and 10mM HEPES (Gibco). MEF cells were grown in DMEM with 15% FBS. On initial receipt, cell lines were expanded and low passage stocks cryopreserved. Cells were regularly tested for mycoplasma by PCR. AZD0156 (cat# S8375) and AZD1390 (cat# S8680) were purchased from Selleck Chemicals and reconstituted in 100% ethanol.

### 2.2 Clonogenic survival

Cells were treated with AZD0156 or AZD1390 for 1 hour and then irradiated with 0-6 Gy. IR was delivered using a PXI X-RAD 225 XL irradiator at maximal dose rate of 1.87 Gy/min with a beam conditioning aluminum filter (Precision X-ray Irradiation). Following IR, cells were counted, re-plated at varying cell numbers to ensure adequate colony formation. 7-14 days after seeding, formed colonies were methanol fixed, stained with Giemsa, and counted. Surviving fraction was calculated as fraction of irradiated cells surviving in relation to non-irradiated cells.

### 2.3 Cytotoxicity assay

Human and murine cancer cell lines or normal tissue MEFs were plated in 96 well plates and exposed to an ATM inhibitor concentration range of 0 – 1000 nM for 72 hrs. Alamar Blue was added to the cells and allowed to incubate for 2-4 hours at 37°C. Plates were analyzed using a plate reader with fluorescence measured at 560 nm. Fractional survival was normalized to untreated control cell fluorescent values.

### 2.4 γH2AX immunostaining

Cells were grown on glass coverslips, treated with 10 nM ATM inhibitor for 1 hour and then irradiated with 2 Gy. Cells were fixed at 1, 2 and 24 hours after irradiation with 4% paraformaldehyde, permeabilized with 0.25% Triton X-100 and stained with antibody to γH2AX diluted to 1:750 (Millipore, cat# 05-636). Nuclei were stained with 4’, 6-diamidino-2-phenylindole (DAPI). Foci were counted over multiple high-power fields per group.

### 2.5 Synthesis of ACPP-AZD0156

Synthetic scheme and chemical structures of cRGD-ACPP-AZD0156 are shown in **Supplementary Fig. S1**. AZD0156 was first attached to maleimidocaproyl-valine-citrulline-p-aminobenzyl hydroxyl (MC-VC-PAB-OH) to create MC-VC-PAB-AZD0156 **Supplementary Fig. S1**. MC-VC-PAB-OH (6.6 mg, 11.5 µmol; Ambeed) was suspended in dry DMF (50 µl) under dry nitrogen with stirring at room temperature in a capped plastic vial. Three additions of thionyl chloride (6.5 µL, 2 M solution in DCM, 13 µmol) were added over 2 hours and volatile solvents were removed by a stream of nitrogen to give a pale-yellow hazy solution. AZD0156 (3.79 mg, 8.21 µmol) dissolved in dry DMF was added followed by DIEA (5 µL, 29 µmol) and the hazy solution was stirred overnight. Additional DIEA (5 µL) was added, a hole made in vial cap and stirred overnight. After LC-MS showed the desired product, acetic acid (20 µL) was added, and the product separated by semi-prep HPLC and lyophilized to a white solid as the TFA salt with a yield of 73% (6.74 mg, 5.96 µmol). ES-MS (m/z) [M^+^] found 1015.2 calculate for C_54_H_70_N_11_O_9_^+^, 1016.5, **Supplementary Fig. S1**.

To synthesize ACPP-AZD0156, H_2_N-peg8-e_9_-oPLGC(Me)AG-r_9_-c-CONH_2_ where lower case letters refers to D-amino acids, peg8 refers to H_2_N-PEG8-propionic acid, o-denotes 5-amino-3-oxopentanoyl (a short hydrophilic spacer), C(Me) denotes for S-methylcysteine and the CONH_2_ indicates C-terminal amide (10·TFA salt, 10.43 mg, 2.14 µmol) dissolved in dry DMSO (100 µL) was mixed with MC-VC-PAB-AZD0156 (214 µL, 10 mM solution in DMSO, 2.14 µmol) and NMM (2.35 µL, 21.4 µmol) and kept at room temperature. LC-MS indicated complete reaction after 30 min to give a single product that was used without further purification. ES-MS (m/z) [M^+^] found 947.9 (M^+^+4H^+^), 1185.6 (M^+^+3H^+^), deconvolved to 4738.2, calculated for C_201_H_329_N_66_O_63_S_2_^+^, 4739.4. A solution of 6-maleimidocaproic acid N-hydroxysuccinimide ester (Sigma-Aldrich; 21.4 µl of 100 mM in dry DMSO, 2.0 µmol) was added to the reaction mixture and kept at room temperature for 4 days until LC-MS showed reaction was complete. ES-MS (m/z) [M]^+^ found 986.9 (M^+^+4H^+^), 1233.8 (M^+^+2H^+^), deconvolved to 4930.9 calculated for C_211_H_34_0N_67_O_66_S_2_^+^, 4932.47. Cyclo(RGD)fC (Peptides International, 1.4 mg, 2.4 µmol) dissolved in dry DMSO (100 µL) was added and mixed. LC-MS indicated complete reaction after 30 min to yield the final product, and the reaction was quenched with acetic acid (50 µL), separated by HPLC and lyophilized to give cyclo(RGD)fc-MC-HN-peg8-e_9_-oPLGC(Me)AG-r_9_-c-(MC-VC-PAB-AZD0156)-CONH_2_ as a white powder with a Yield of 7.36 mg (52%). ES-MS (m/z) [M^+^] calculated for C_235_H_374_N_75_O_73_S_3_^+^, 5510.7 found 1102.8 (M^+^+4H^+^), 1378.8 (M^+^+3H^+^), deconvolved to 5509.1, **Supplementary Fig. S2**. Peptide product was lyophilized and stored as a powder at −20°C.

### 2.6 Flow Cytometry analysis

All animal work was done in compliance with the University of California San Diego Institutional Animal Care and Use Committee, protocol# S15290. Mice were housed in individually ventilated and micro-isolator cages supplied with acidified water and 5053 irradiated PicoLab Rodent Diet 20 feed. Temperature for laboratory mice in housing was 18-23°C with humidity 40-60%. Housing room was maintained on a 12-hour light/dark cycle. B16 tumors established in 6-week-old female C57BL/6 albino mice were treated with free AZD0156, ACPP-AZD0156 and IR. Tumors were harvested as indicated in **Figure Legends**, minced, and re-suspended in FBS-free DMEM media supplemented with components of MACs tumor dissociation kit. Tissues were incubated for 15 minutes at 37°C and mechanically digested using the gentle MACs Octo Dissociator. Tissue suspensions were washed with fresh media and passed through a 100-µm strainer. Samples were washed with PBS and immediately processed for live/dead cell discrimination using BD LIVE/DEAD™ Fixable Blue Dead Cell Stain Kit. Cells were washed and stained for surface markers for 30 minutes at 4° C. Intracellular staining was performed using the eBioscience FOXP3/Transcription Factor Buffer Set from Invitrogen and stained with intracellular antibodies. Antibodies were purchased from BioLegend and included CD45 (30-F11, 1:100), CD8a (53-6.7, 1:100), CD4 (RM4-4, 1:400), and NK1.1 (PK136, 1:100). All antibodies were validated by the supplier. All flow cytometry data acquisition was done using BD LSRFortessa and analyzed using FlowJo software. Immune cells were identified by the following characteristics: cytotoxic T cells (CD45+Thy1.2+CD8+), helper T cells (CD45+Thy1.2+CD4+), NK cells (CD45+Thy1.2-NK1.1+). Characterized CD8+ T-cells were further gated for granzyme-B (GB11)

### 2.7 NanoString analysis

B16 tumors were established as above in 6-week-old female C57BL/6 albino mice. Mice were then treated with ACPP-AZD0156 and IR. Tumors were harvested as indicated in **Figure Legends**. For harvesting, tumors were snap frozen in liquid nitrogen. RNA was isolated from tumor tissues and comprehensive gene expression immune profiling was analyzed using the mouse NanoString PanCancer IO 360 Panel platform. The Advanced Analysis module of the nSolver software was used to analyze genes associated with listed immune cells in tumors and given a Z-score (n = 3 mice per group).

### 2.8 In vivo subcutaneous tumor therapy experiments

B16 tumor cells were injected subcutaneously into the hindlimbs of 6-week-old female C57BL/6 albino mice in 100 µl of 1:1 Growth Factor Reduced Matrigel (BD) and PBS solution. Mice were injected with free AZD0156 or ACPP-AZD0156 on days 3, 4 and 5 post-implantation. Fractionated 5 Gy IR was delivered focally to the tumor bearing hindlimbs B16 tumors on days 3, 4 and 5 two hours after AZD0156 injection using a PXI X-RAD 225 XL irradiator at maximal dose rate of 3 Gy/min with a beam conditioning copper filter (Precision X-ray Irradiation). For CD8 depletion studies, mice were intraperitoneally (i.p.) injected with anti-CD8 (BioXCell #BE0117) at 10 mg/kg on days −6, −5 and −4 tumor cell implant and then weekly starting on day after tumor cells were injected or control antibody. For anti-PD-1 therapy, mice were i.p. injected with 200 μg anti-PD-1 antibody (BioXCell #BE0033) or control antibody on days 3, 6 and 11 after tumor cell implantation. Tumor volumes were measured and calculated using the formula as ½ * Length * Width^2^. Mice were sacrificed when tumor volume reached > 1500 mm^3^, maximal tumor volume approved on our protocol for subcutaneous tumors.

### 2.9 Statistical Analysis

Unpaired 2-sided t test or ordinary one-way ANOVA were performed for quantitative cell culture clonogenic survival, γH2AX foci, flow cytometry and end of study tumor volume responses. Survival curves were analyzed Log-rank (Mantel-Cox). All statistical analyses were performed using Prism software (GraphPad).

## 3. Results

### 3.1 Tumor targeted cell penetrating peptides

To address the challenges of tumor specific radiosensitization, we developed molecularly guided activatable cell penetrating peptides (ACPPs) that redirect extracellular tumor proteases for oncologic applications[42-45]. The ACPP drug delivery platform is a scaffold designed to cloak cell penetrating peptides that is then specifically “activated” by matrix metalloproteinase (MMP-2 and MMP-9) that are overexpressed within tumors [46-48]. ACPPs comprise a polycationic cell penetrating domain (nine repeats of d-arginine, r9) electrostatically masked by a polyanionic domain (nine repeats of d-glutamic acid, e9) through MMP-2/9 sensitive PLGC(Me)AG peptide linker. MMP-2/9 cleavage of ACPP results in release of the r9 cell penetrating peptide which can electrostatically adhere to cancer cell membranes and internalize in vesicles. These vesicles containing ACPP then fuse with the endosomes followed by its trafficking to the lysosomes (**Figure 1A**). Using identical drug-linker chemistry clinically validated with antibody drug conjugates (ADCs), drugs can be conjugated to the cell penetrating peptide moiety of ACPPs[49, 50]. To validate MMP-2 and MMP-9 is overexpressed in tumors compared to normal tissues, we established syngeneic MC38 tumors in immune-competent mice and measured gelatinase activity in tumors compared to adjacent normal tissue (i.e. peri-tumoral) muscle as well as vital organs (i.e. heart, lung, liver, and kidney) using gelatin zymography. Tumors showed increased MMP-2 and MMP-9 activity whereas all normal tissues tested had minimal MMP activity including peri-tumoral muscle (**Figure 1B**). These results support the rationale for tumor specific drug delivery of the ACPP platform and its use to study how targeted DDR inhibitors can modulate the irradiated tumor immune microenvironment.

**Figure 1:**
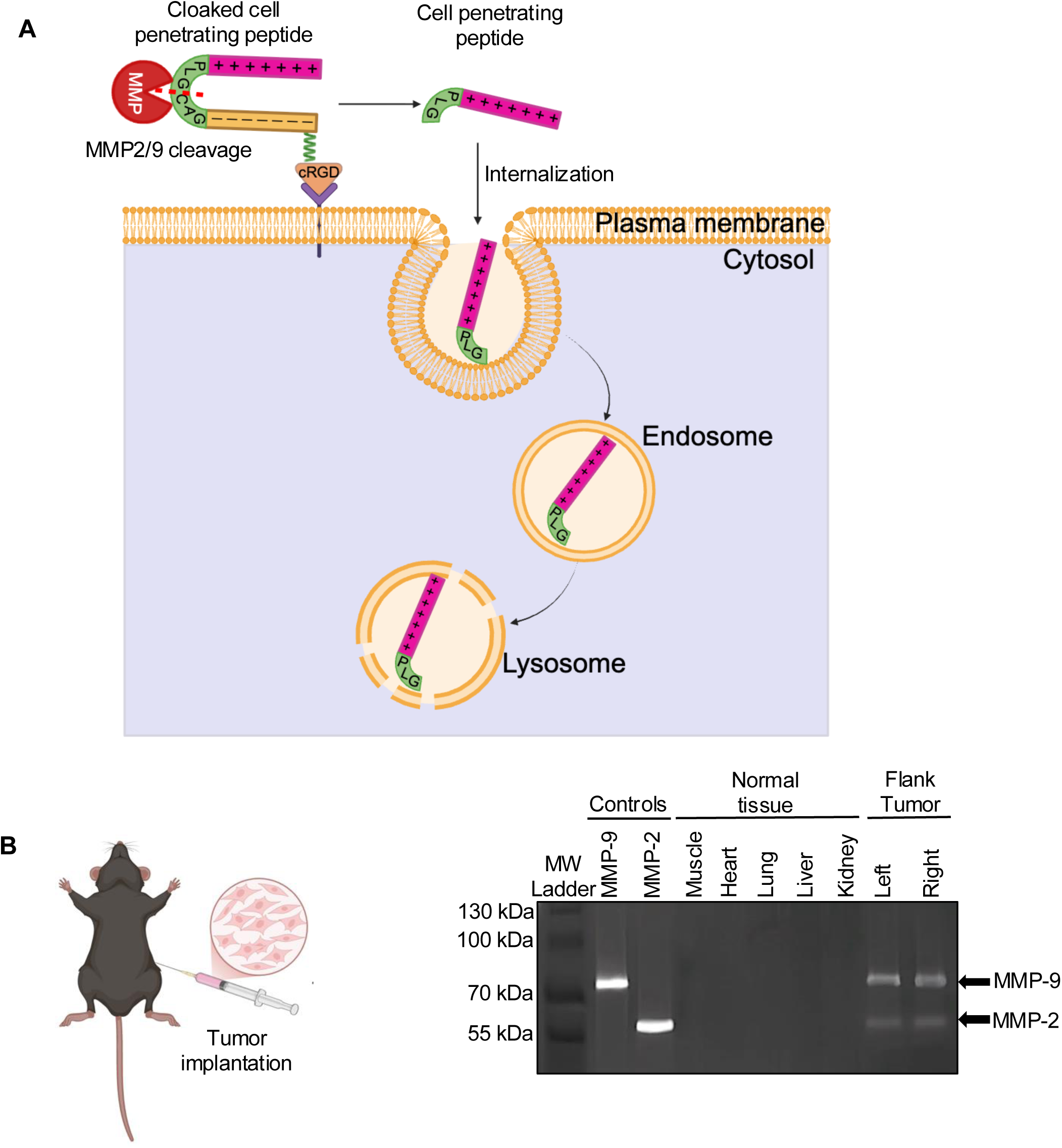
ACPP directed tumor-targeting in syngeneic mouse model. **A)** The ACPP construct consists of a polycationic r9 (d-arginine) domain masked by a polyanionic e9 (d-glutamate) domain connected via a matrix metalloproteinase (MMP)-sensitive linker [PLGC(Me)AG] and an integrin-targeting cRGD motif. Overexpressed MMP2/9 in tumor microenvironment cleave the ACPP linker that allows its internalization into vesicles at the plasma membrane. The vesicles containing ACPP construct then fuse with the endosomes followed by ACPP trafficking to the lysosomes. **B)** MC38 tumors are implanted into mice hindlimb. Gelatin zymography of MMP2/9 activity from recombinant MMP controls, normal murine tissues (muscle, heart, lung, liver and kidney), and lysates from mice with established syngeneic MC38 tumors. Red arrows mark MMP-2 and MMP-9 activity. Molecular weight (MW) ladder shown in left lane.

### 3.2 Non-specific radiosensitization by ATM inhibitors

ATM plays a central role in coordinating DNA damage response following IR induced DNA damage and subsequent repair of DNA double strand breaks[19, 20]. Recently, potent next generation ATMi have been developed that include AZD0156 and AZD1390[22, 23]. Importantly, both AZD0156 and AZD1390 are chemically amenable to peptide-drug conjugation using validated ADC drug linker chemistry[51, 52]. We first evaluated the radiosensitization potency of AZD0156 and AZD1390 across a panel of human (A549, LN229, CAL27) and murine (LL2, MC38, B16) cancer cell lines using the surviving fraction at 2 Gy clonogenic assay (SF2). The SF2 assay uses a clinically relevant 2 Gy dose of IR that is routinely given to patients undergoing conventionally fractionated radiotherapy. Briefly, cells were treated with 10 nM of ATMi for 1 hour followed by 2 Gy irradiation. Cells were then replated in drug-free media and cell viability measured by colony growth formation 7-14 days post-IR. Both AZD0156 and AZD1390 significantly increased radiation cell kill across all cell lines tested, with AZD0156 consistently demonstrating more potent radiosensitization than AZD1390 (**Figure 2A)**. We then tested the radiosensitization potency of AZD0156 and AZD1390 as a function of IR dose. Murine cancer cell lines were treated with 10 nM ATMi for 1 hour followed by IR doses of 1, 2, 4 and 6 Gy. Again, both ATM inhibitors radiosensitized, with AZD0156 showing increased potency compared to AZD1390 (**Figure 2B**). Given that AZD0156 was the more potent ATMi radiosensitizer, we focused the remainder of our studies on AZD0156.

**Figure 2:**
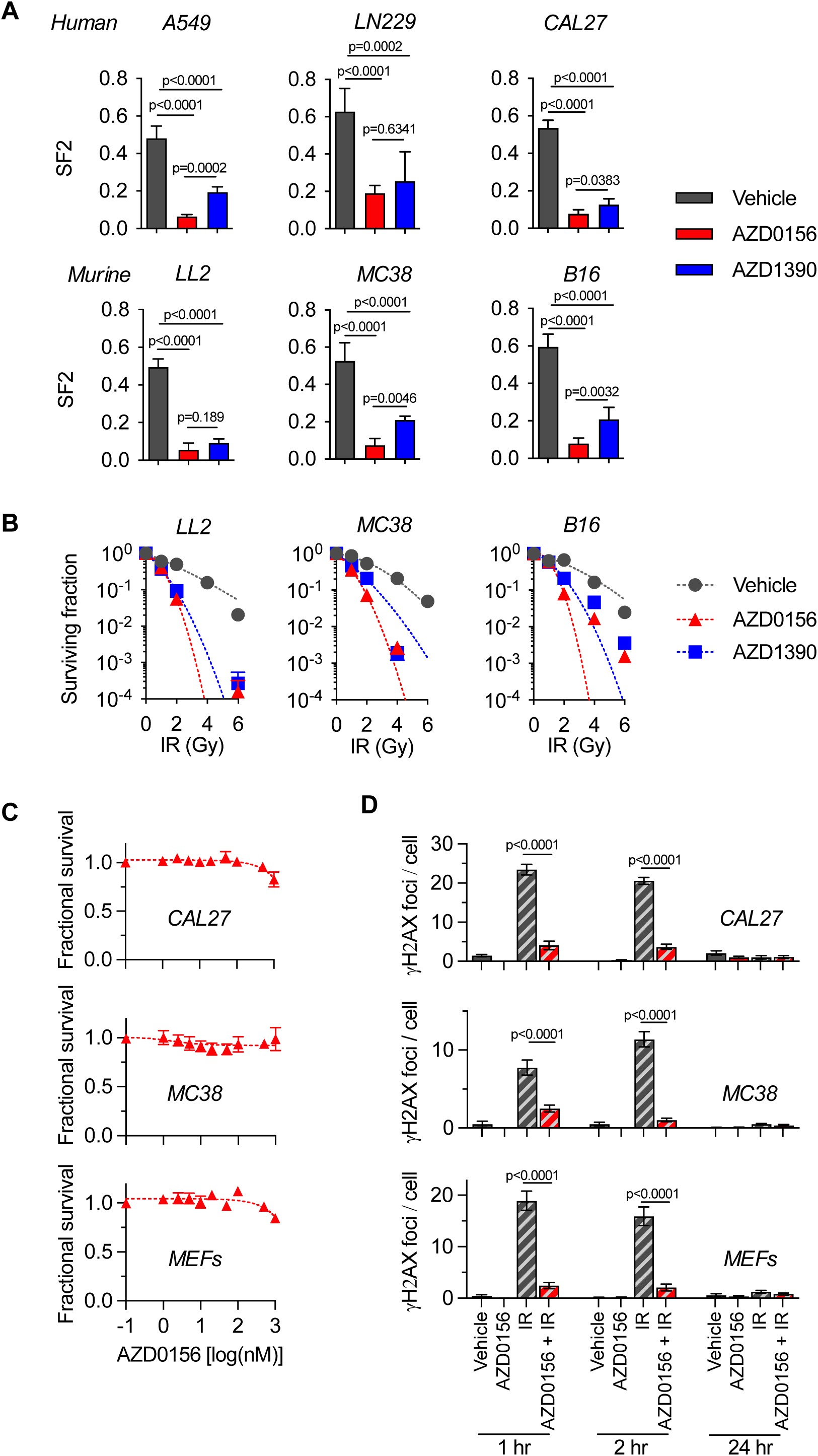
ATM inhibitors non-selectively radiosensitize. **A)** Surviving fraction at 2 Gy (SF2) of human (top panel) and murine (lower panel) cancer cells that were treated with 10 nM of AZD0156 or AZD1390 and irradiated with 2 Gy. Cells were fixed 2 hours post-irradiation and analyzed for γH2AX foci formation by immunofluorescence, n>60 cells/condition. **B)** Clonogenic survival of murine cancer cells treated with 10 nM of ATM inhibitors and irradiated with 0-6 Gy. Cell viability normalized to non-irradiated cells for each drug and plotted as mean fractional survival ± SEM, n=6. Non-linear regression curve fit shown for each drug. **C)** Human and murine cancer cells or normal tissue murine embryonic fibroblasts (MEFs) were exposed to a dose range of AZD0156 for 72 hours. Cell viability was measured by Alamar Blue assay, normalized to control untreated cells and plotted as mean fractional survival ± SD, n=6. **D)** Cancer cells (CAL27 and MC38) or normal tissue murine embryonic fibroblasts (MEFs) were treated with AZD0156 and irradiated with 2 Gy. Cells were fixed at 1-, 2- and 24-hours post-irradiation and analyzed by immunofluorescence for γH2AX foci formation. Statistical significances calculated using one-way ANOVA with Turkey’s multiple comparisons test.

Although chemotherapies are given concurrently with radiotherapy to also radiosensitize, they have systemic toxicities that prevent further treatment intensification. Based on their mechanism of action, ATMi provide a non-cytotoxic approach to radiosensitize tumors. To confirm this, we measured the intrinsic cytotoxicity of AZD0156 in human (CAL27) and murine (MC38) cancer cells along with non-transformed murine embryonic fibroblasts (MEFs). Notably, AZD0156 showed minimal cytotoxicity across a broad dose range (0-1000 nM) in both cancer cells and non-cancerous cells (**Figure 2C**). Finally, we tested the radiosensitization activity of AZD0156 in either cancer or normal cells based on DNA damage response signaling. We measured γH2AX (pS139 histone H2A) foci formation that is a validated marker for DNA double-stand break repair. While AZD0156 blocked IR-induced γH2AX foci accumulation in cancer cells, it was equally active in irradiated non-cancerous MEFs, indicating non-selective blockade of DNA repair (**Figure 2D**). Thus, although ATMi are effective radiosensitizers, free-drug ATMi indiscriminately radiosensitize all exposed cells within the irradiated target volume including normal cells, highlighting the need for tumor-restricted delivery to improve their therapeutic index when given with radiotherapy.

### 3.3 Tumor-targeted ACPP-ATMi synthesis

To synthesize a radiosensitizing tumor-targeted cell penetrating peptide-ATMi conjugate conjugate, we coupled AZD0156 to the ACPP scaffold using linker chemistry of the clinically validated maleimidocaproyl-valine-citrulline-p-aminobenzylcarbamate (MC-VC-PABC) linker that is part of FDA-approved MMAE containing ADCs (**Figure 3A**)[51-53]. An advantage of the MC-VC-PABC linker is that following drug conjugate intracellular uptake into endolysosomes, the intervening valine-citrulline dipeptide of the linker is cleaved by lysosomal cathepsins resulting in self-immolative loss of p-iminoquinone methide with CO2 and predictable release of the free drug molecule. AZD0156 was chosen as our lead ATMi for ACPP conjugation since it had increased radiosensitizing potency compared to AZD1390 (**Figure 2**). Due to the absence of a primary or secondary amine on AZD0156, we derivatized its tertiary amine to a quaternary ammonium group by alkylation with maleimidocaproyl-valine-citrulline-p-aminobenzylchloride (MC-VC-PAB-Cl) to attach the self-immolative MC-VC-PAB moiety[54, 55], yielding MC-VC-PAB-AZD0156 (**Figure 3B, S1**). The charged quaternary ammonium group also has the added advantage of increasing the hydrophilicity of the linker-drug. The intermediate and final ACPP-AZD0156 conjugate were purified by reverse-phase HPLC and confirmed by mass spectrometry (**Figure 3C-D, S2**).

**Figure 3:**
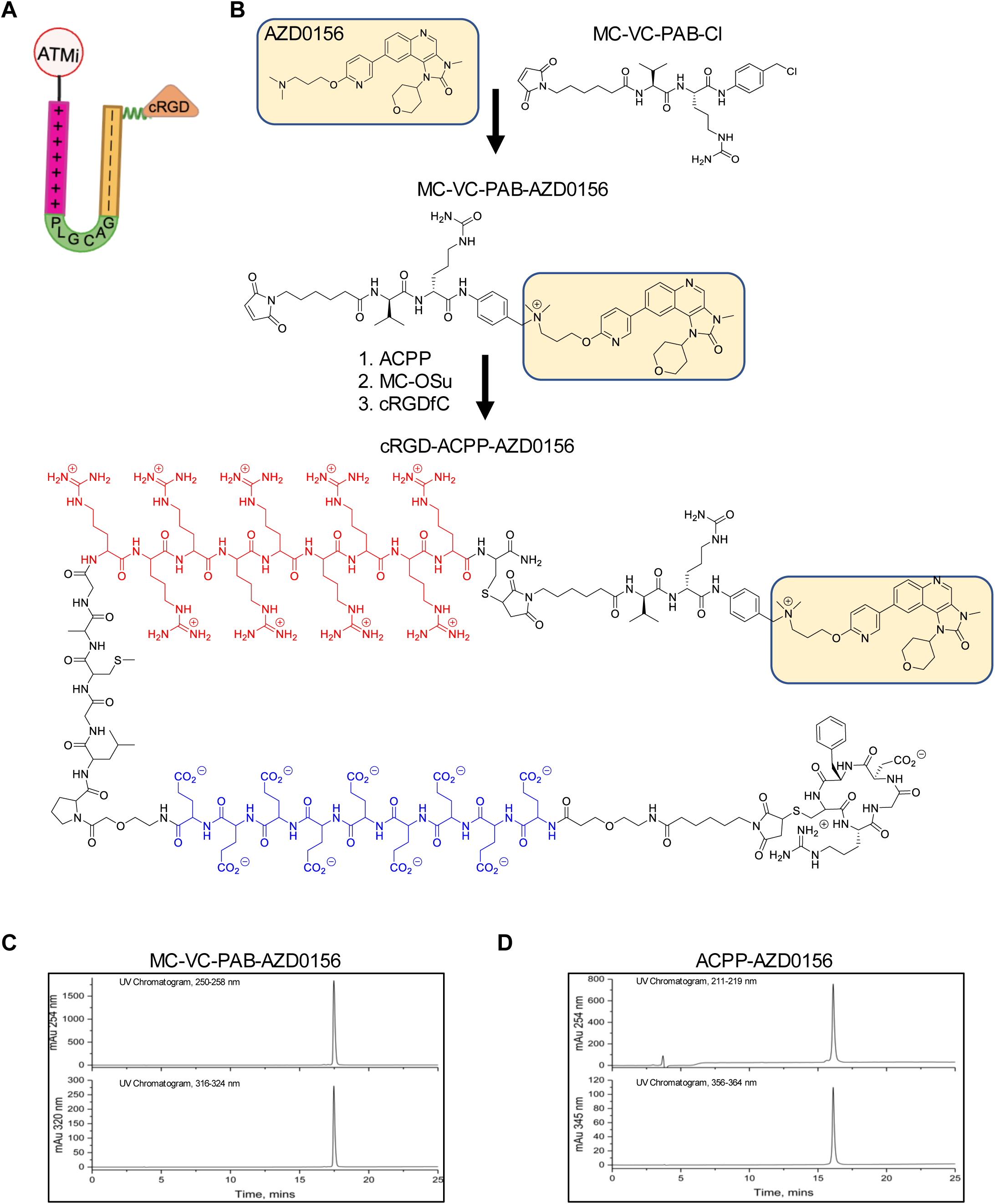
Synthesis of ACPP-AZD0156 conjugate. **A)** Schematic illustration of ACPP-AZD0156 conjugate. The construct consists of a polycationic cell penetrating peptide (+), autoinhibitory polyanionic peptide(-), a matrix metalloproteinase sensitive PLGC(Me)AG peptide linker, cycloRGD, and ATMi drug payload. **B)** Stepwise conjugation of ATMi to ACPP. AZD0156 was first derivatized using a maleimidocarpoyl-valine-citrulline-p-aminobenzyl (MC-VC-PAB) cathepsin-B-cleavable linker to form the MC-VC-PAB-AZD0156 intermediate. The polycationic cell-penetrating peptide domain (r9, shown in red) and the polyanionic autoinhibitory domain (e9, shown in blue) were linked via an MMP2/9-sensitive peptide linker, followed by maleimide coupling of MC-VC-PAB-AZD0156 to the r9 domain of ACPP and cRGD attachment to the e9 domain for α_v_ý_3_ integrin pre-targeting. Reverse-phase HPLC analyses of **(C)** MC-VC-PAB-AZD0156 and **(D)** ACPP-AZD0156. Elution conditions were 5-50% acetonitrile in water with 0.05% TFA over 20 minutes, then to 90% over 10 minutes at 1ml/min using a Luna(2) C18 column, monitored at 215 and 360 nm.

### 3.4 Immune activation by tumor-targeted ACPP-AZD0156 radiosenstization

Since ATM inhibition can potentiate type I interferon signaling and enhance anti-antitumor immune response in pre-clinical models[38, 39, 41, 56], we next determined how tumor-restricted targeted delivery of ATMi modulated the irradiated tumor immune microenvironment. For a non-biased approach, we first carried out transcriptomic profiling (NanoString) on B16 tumors treated with ACPP-AZD0156, IR, or the combination of both. Starting on day 10 post tumor cell implantation, mice were treated for 3 consecutive days with ACPP-AZD0156 and IR. ACPP-AZD0156 was intravenously injected at a dose of 10 nmol (3.3 mg/kg) per day (30 nmol total). IR was focally delivered to the tumor bearing hindlimb, 5 Gy per day (15 Gy total). On each treatment day, ACPP-AZD0156 was injected first followed by irradiation, to understand the mechanism of action of ATMi radiosensitization. Tumors were then harvested on day 18 (8 days after initiation of therapy) and RNA was isolated (**Figure 4A**). RNA samples were subjected to NanoString IO panel. Principal component analysis showed that the combination treatment of IR with ACPP-AZD0156 produced a transcriptionally distinct cluster compared to control (untreated) or either monotherapy (IR alone or ACPP-AZD0156 alone) (**Figure S3A**), indicating that targeted ATMi radiosensitization produced a qualitatively unique biological response. Volcano plots (**Figure 4B)** and Gene Ontology network analysis showed that ACPP-AZD0156 alone induced modest differential gene expression that primarily involved cytokine activation and antiviral/type I interferon pathways (**Figure 4C**, red curves). IR alone upregulated genes involved in cytokine signaling, innate immune activation, and antigen presentation (**Figure 4D**, blue curves). In contrast, IR + ACPP-AZD0156 showed a leukocyte network stimulation including genes associated with cytotoxic T-cell activation, immune synapse formation, Fc-receptor engagement, and antigen-processing machinery (**Figure 4E**, green curves). Differential expression heatmap and pathway enrichment corroborated these findings. The combination therapy of IR + ACPP-AZD0156 enriched unique pathways involved with adaptive immune activation, antigen presentation, dendritic cell priming, leukocyte recruitment, and type I interferon amplification (**Figure S3B, C**). Taken together, these results indicate that tumor-restricted ATM inhibition does not merely augment radiation-induced immune engagement but instead synergizes to induce a broader immune-stimulatory transcriptional program that includes a leukocyte activation network not triggered by IR or ACPP-AZD0156 alone.

**Figure 4.**
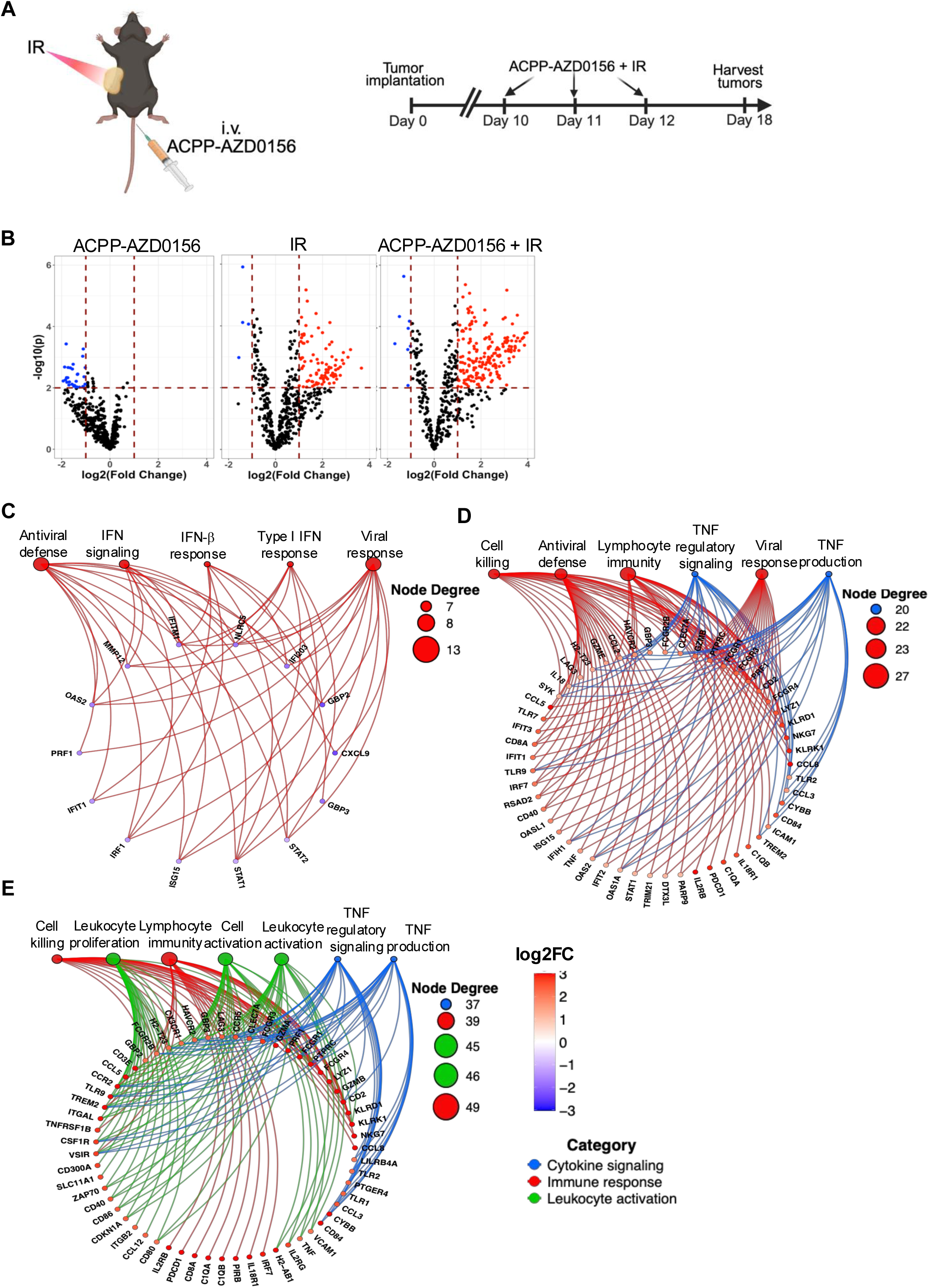
Gene expression alterations induced by tumor-targeted radiosensitization. **A**) Schematic illustration of treatment to the mice. Mice with B16 tumors were treated with ACPP-AZD0156, IR alone, or ACPP-AZD0156 + IR starting on day 10 post tumor cell implantation. Tumors were harvested on day 18 and analyzed using NanoString nCounter PanCancer IO 360 panel. **B**) Volcano plots for gene expression of ACPP-AZD0156 (left), IR alone (middle), and ACPP-AZD0156 + IR (right) treated B16 tumors vs control untreated samples, n=3. Dashed lines represent thresholds for significance in terms of p-value (horizontal lines) and log2 fold change (vertical lines). Red dots indicate significantly upregulated genes and blue dots indicate significantly downregulated genes in treated cells. **C-E**) Gene Ontology network analyses of significantly altered genes from ACPP-AZD0156 (C), IR alone (D), and combination treatment (E). Nodes represent either GO biological pathways or significantly differentially expressed genes, and curves indicate gene association within a given GO term. Node size corresponds to node degree, defined as the number of genes associated with a pathway. Source data in Source Data file.

### 3.5 ACPP-ATMi radiosensitization modulates immune composition in vivo

Given the induction of immune-activating transcriptional programs, especially the leukocyte activation network, we next tested if these transcriptional changes translated into altering the tumor immune infiltrating cells in tumors from treated mice. To test this, syngeneic murine B16 tumors were once again established in immune-competent mice as mentioned above in **Figure 4A**. Tumors were then harvested on day 18 (8 days after initiation of therapy) and tumor immune infiltration was assessed by flow cytometry. Tumors from untreated control mice and ACPP-AZD0156 showed minimal CD8+ T-cell infiltration, 2.6% and 1.3% of total live cells respectively (**Figure 5A, B**). IR alone induced significant CD8+ T-cell infiltration (32.3%). Importantly, the combination of IR + ACPP-AZD0156 further increased CD8+ T-cell tumor infiltration to 58%, significantly exceeding IR alone. For CD4+ T-cells, IR again increased this T-cell population from 1.1% (untreated tumors) to 6.3 %. Unlike the effects on CD8 T-cell tumor infiltration, ACPP-AZD0156 radiosensitization did not significantly increase CD4+ T-cells beyond levels induced by IR, indicating its adaptive immune effects are primarily CD8-dominant. For NK cells, IR increased this immune cell population compared to untreated tumors but decreased when ACPP-AZD0156 was added, consistent with a shift from innate to adaptive immunity as seen by our transcriptional profiling (**Figure 4E**). Moreover, additional arms of the immune system including dendritic cells, macrophages and neutrophils also increased in response to ACPP-AZD0156 radiosensitization compared to IR or ACPP-AZD0156 monotherapies (**Figure 5C**). Taken together, comprehensively interrogating tumor immune infiltration in an unbiased fashion by gene expression analysis corroborated flow cytometry studies that ACPP-AZD0156 radiosensitization not only enhances the magnitude of CD8+ T-cell infiltration but also qualitatively reshaped the immune landscape which could prime the tumor microenvironment for adaptive immune responses to synergize with immune checkpoint therapies.

**Figure 5.**
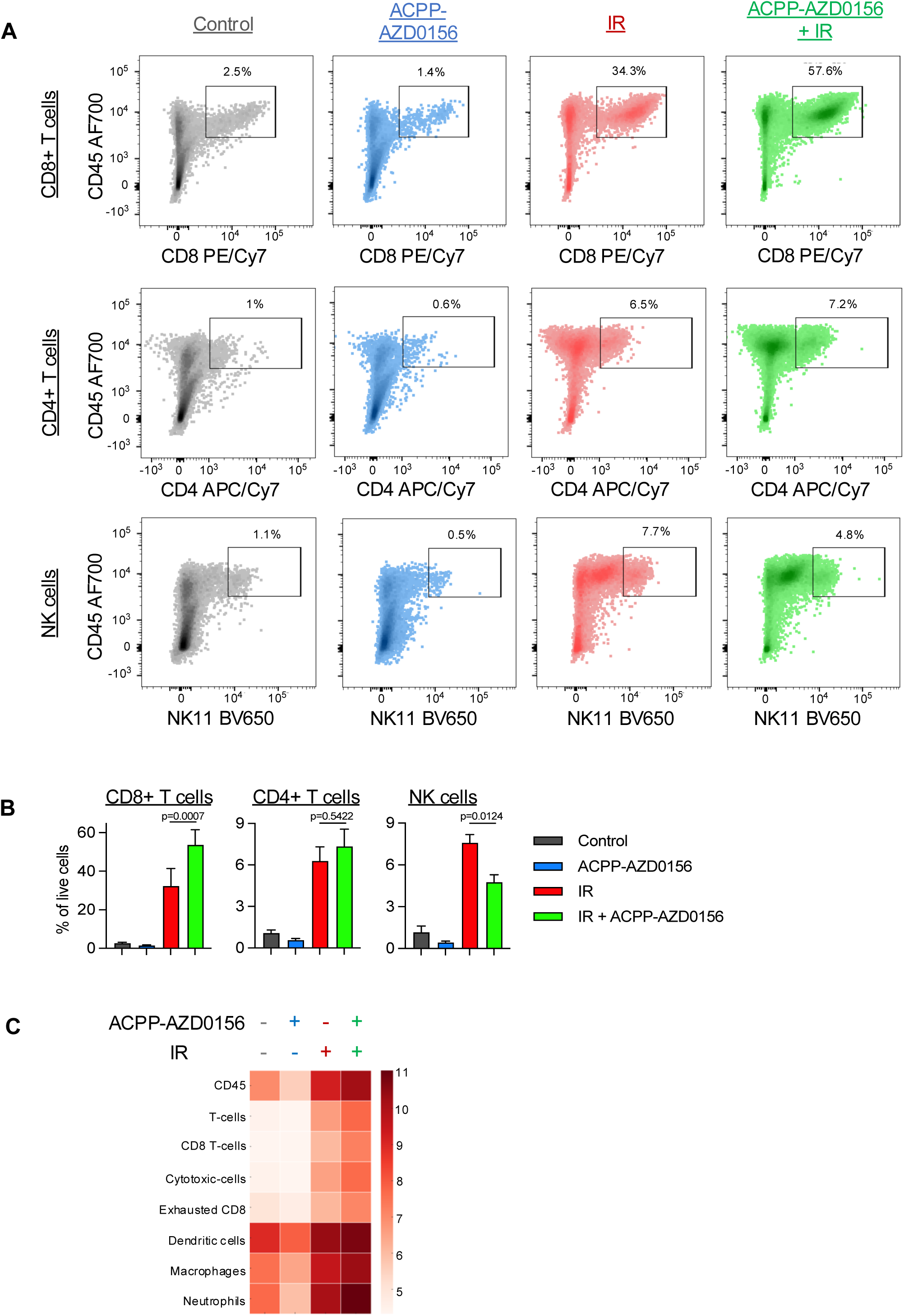
ACPP-ATMi radiosensitization stimulates tumor immune-infiltration. **A)** Mice with B16 tumors were treated as mentioned in Figure 4A. Tumors were harvested on day 18 and CD8+, CD4+ and NK tumor immune infiltration were characterized by flow cytometry and **(B)** plotted as mean ± SEM. Statistical significances calculated using one-way ANOVA with Turkey’s multiple comparisons test. **C)** Mice with B16 tumors were treated as in Figure 5A. Tumors were harvested on day 18 and analyzed using NanoString nCounter PanCancer Mouse Immune Profiling panel. Heatmap depicts Z-scored immune cell-type signature scores across treatment groups.

### 3.6 ACPP-ATMi radiosensitization potentiates immunotherapy

Next, we asked whether immune remodeling could achieve durable tumor control. Murine B16 tumor cells were implanted into immune-competent mice. We used the same treatment scheduling as for tumor immune profiling studies. At 9 days post-tumor cell implantation, mice were i.v. injected with 10 nmol of either free AZD0156 (0.23 mg/kg) or ACPP-AZD0156 (3.3 mg/kg) for 3 consecutive days (30 nmol total). IR was given on each of the 3 days of drug injection with 5 Gy fractions given focally to the tumor bearing hindlimb (15 Gy total). Free AZD0156 or ACPP-AZD0156 monotherapies did not significantly decrease tumor growth compared to untreated controls (**Figure 6A**). Notably, combination treatment with IR + ACPP-AZD0156 significantly reduced tumor growth compared with IR alone or IR combined with free AZD0156 despite equivalent dosing, highlighting the value of spatial drug targeting. In addition, flow cytometric analysis showed that ACPP-AZD0156 combined with IR enhances activated CD8+ T cells more effectively than free AZD0156 when combined with IR (**Figure 6B**). Finally, cytotoxic T cell function was enhanced as measured by the significant increase in Granzyme B expression in the IR + ACPP-AZD0156 combination group, compared to IR + free AZD0156, p=0.0684.

**Figure 6.**
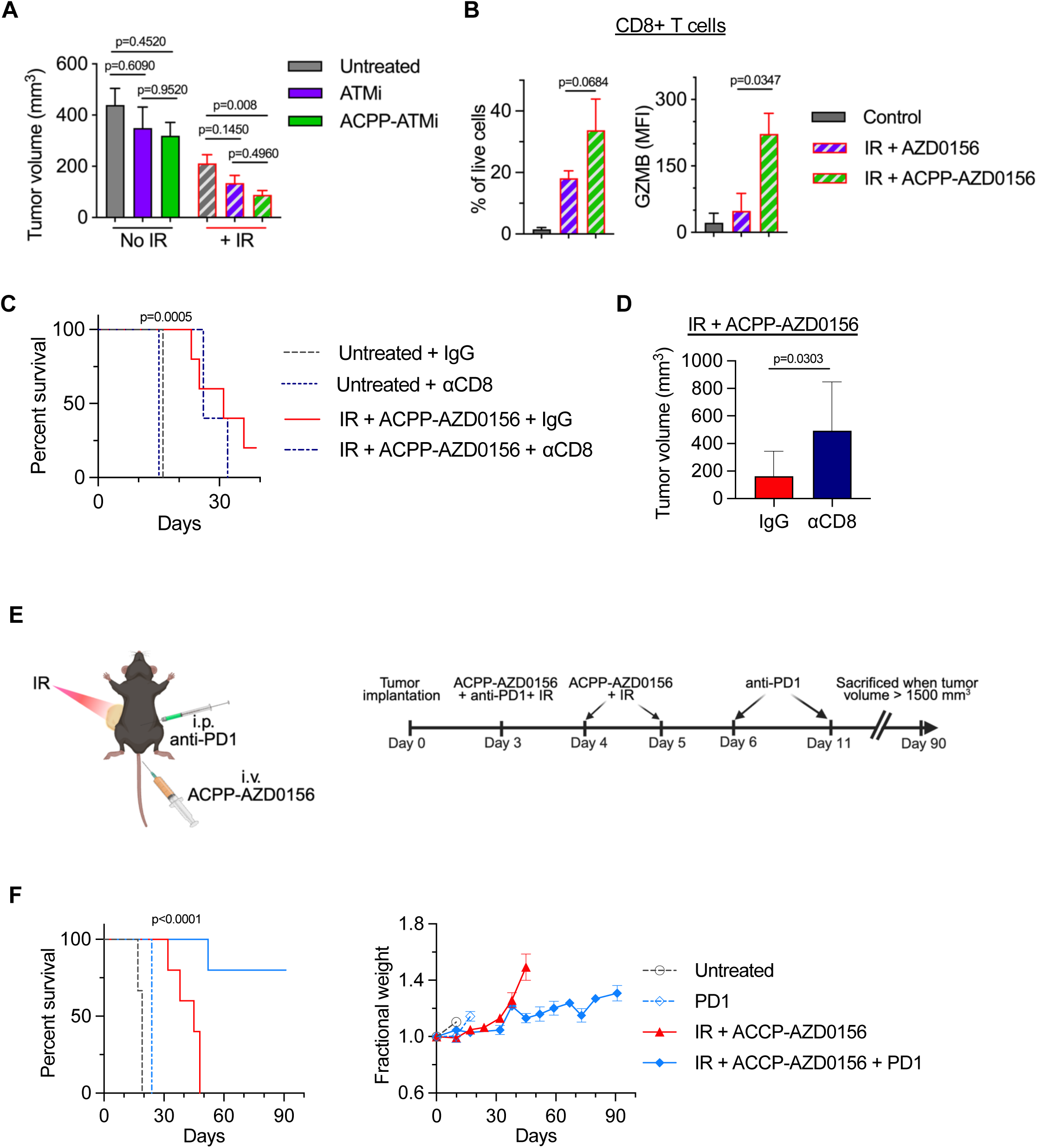
ACPP-AZD0156 potentiates immune-checkpoint tumor control. **A)** Mice with B16 tumors were treated with combination of free AZD0156 or ACPP-AZD016 and IR on day 9 post tumor cell implantation and day 15 tumor volume (mean ± SEM) was plotted, n=10-14. Statistical significances calculated using one-way ANOVA with Turkey’s multiple comparisons test**. B)** Tumors from mice treated as in Figure 4A were harvested on day 18 and CD8+ T-cell infiltration and granzyme B expression was measured by flow cytometry and plotted as mean ± SEM, n=3 (vehicle), n=4 (all other groups). Statistical significance was calculated using two tailed t-tests. **C)** Mice with B16 tumors treated with ACPP-AZD0156 and IR starting on day 3 post tumor cell implantation. Anti-CD8 antibody administration was initiated 6 days prior to tumor cell implantation. Kaplan-Meier survival curves for mice were plotted and statistical significance calculated using log-rank (Mantel-Cox) test, n=3 (Untreated + IgG), n=5 (all other groups). **D)** Tumor volumes at day 25 in immune competent (IgG) or CD8-depleted mice treated with IR + AZD0156 as in Figure 5A, plotted as mean ± SD, n=8 (IgG), n=10 (αCD8). **E)** Schematic illustration of mice treated with anti-PD1 immunotherapy. Mice were treated with IR + ACPP-AZD0156 with or without anti-PD1 immunotherapy initiated on day 3 post tumor cell implantation. ACPP-AZD0156 and IR were given on days 3, 4, and 5 post tumor implantation and anti-PD1 antibody was i.p injected on days 3, 6, and 11 after tumor implantation. **F**) Survival analysis of Statistical significance calculated using Log-rank (Mantel-Cox) test, n=3 (Untreated, αPD1 groups), n=5 (IR + ACPP-AZD0156 ± αPD1). Weekly fractional mouse body weight plotted, normalized to 1 at start of study (day 0).

We then investigated the role of adaptive immune responses on ACPP-AZD0156 radiosensitized tumor control. CD8 depletion abrogated tumor control in the combination arm (**Figure 6C, D**). By day 25 (the last day where all mice survived in IR + ACPP-AZD0156 groups), the mean tumor volume of IR + ACPP-AZD0156 treated tumors was significantly smaller 163.5 mm^3^ in immune-competent mice compared to 492.9 mm^3^ in mice where CD8+ T cells were depleted, establishing that therapeutic efficacy of ACPP-ATMi radiosensitization was CD8+ T-cell dependent. Since tumor control by ACPP-AZ0156 radiosensitization was mediated by the adaptive arm of the immune system, we then tested whether tumor-targeted radiosensitization could synergize with immune checkpoint inhibitor therapy (**Figure 6E**). While tumors eventually grew out in all mice treated with IR + ACPP-AZD0156, the addition of anti-PD-1 therapy resulted in durable tumor control and long-term survival in 80% of treated mice (4 of 5 mice). Since in our model systems tumors were grown in the bilateral flanks (2 tumor per mouse), 8 of 10 tumors in mice treated with IR + ACPP-AZD0156 + anti-PD1 antibody regressed completely beyond 90 days after implantation in this highly aggressive syngeneic B16 cancer model (**Figure 6F**). From a safety standpoint, mice tolerated this trimodal precision therapy well as assessed by body weight. These data demonstrate that spatial radiosensitization can potentiate immune checkpoint inhibition that results in durable tumor control.

## 4. Discussion

Radiotherapy is a mainstay in the curative treatment of solid tumors, yet its efficacy is limited by tumor resistance and collateral toxicity to surrounding normal tissues[1, 2]. Although cytotoxic chemotherapy, such as cisplatin or taxanes are given with radiotherapy to improve outcomes, their systemic toxicities limit further dose intensification. To address these problems, efforts have focused on molecularly targeted radiosensitizers that specifically inhibit DDR pathways, including ATM, ATR, and DNA-PK to selectively impair repair of IR-induced DNA double strand breaks[13, 39, 41, 57, 58]. Despite preclinical activity, the clinical translation of DDR inhibitors has been challenging. Specifically, clinical trials combining DDR inhibitors with concurrent chemoradiotherapy have caused significant normal tissue toxicities within the irradiated field[25-28]. These data highlight a key limitation of non-targeted DDR radiosensitizers that they indiscriminately radiosensitize both tumor and normal tissues within the irradiated field, thereby negating any improvement to the therapeutic index of radiotherapy. These clinical limitations highlight the need to develop spatially targeted radiosensitizer delivery strategies.

Here, we engineered a tumor-targeted activatable cell-penetrating peptide-ATM inhibitor conjugate (ACPP-AZD0156) that spatially restricts radiosensitization by leveraging tumor-overexpressed matrix metalloproteinases (MMP-2/9) for targeted tumor delivery. The ACPP platform has modular design where a polycationic r9 cell-penetrating peptide is masked by a polyanionic domain through a flexible MMP-cleavable peptide linker[48, 59]. To facilitate clinical translation, we employed the same cathepsin B-cleavable MC-VC-PABC peptide linker used in FDA-approved antibody-drug conjugates (ADCs) that was pioneered in brentuximab vedotin[53, 60]. Upon cleavage in the tumor microenvironment, the unmasked peptide adheres to tumor cell membranes and facilitates internalization and intracellular delivery of the conjugated drug in endolysosomes. The cathepsin-B cleavable MC-VC-PABC linker then enables free drug release following the lysosomal cleavage.

To design our lead ATMi-cell penetrating peptide drug conjugate, we first compared the radiosenstizing potential of two potent ATM inhibitors recently developed, AZD0156 and AZD1390[23, 61]. Across a panel of all human and murine cancer cell lines, AZD0156 consistently was a more potent radiosenstizer compared to AZD1390 and was therefore selected as the payload for ACPP conjugation and for evaluating its anti-tumor effects in an immune competent syngeneic murine model. Unbiased NanoString-based gene expression analysis showed that ACPP-AZD0156 alone induced canonical type I interferon and antiviral response, whereas IR alone activated general inflammatory immune response without interferon-dominant network. Importantly, the combination of IR with ACPP-AZD0156 magnified this response into a broader immune response that uniquely included a leukocyte-activation network absent from either IR or ACPP-AZD0156 monotherapy. This unique network includes cytotoxic effector genes (Granzyme B), chemokines (CCL5, CCL12), antigen-presentation genes (CD86, H2-T23), and inflammatory regulators (IL2RG, TNF), indicating not only enhanced innate immune signaling but also coordinated activation of adaptive immunity. This is consistent with recent studies that DDR inhibitors, including ATM and ATR inhibitors, modulate type I interferon responses and immune infiltration following irradiation[39, 41, 57, 62]. Flow cytometric analysis further validated our transcriptomic findings, showing that ACPP-AZD0156 radiosensitization enhanced intratumoral CD8+ T-cell infiltration and cytotoxic activation marker (Granzyme B) beyond levels seen with IR alone. Importantly, ACPP-AZD0156 was more effective compared to free AZD0156 when combined with IR. Interestingly, no significant changes were observed in CD4+ T-cell populations, and NK cell infiltration was modestly reduced between the combination treatment and IR alone, indicating that the anti-tumor effects were not driven by helper T cell modulation and suggesting a shift toward a CD8-dominant adaptive immune response. These results suggest that targeted ATM inhibition may not enhance innate immunity but rather recruits cytotoxic T-cell responses into irradiated tumors. In support of this, the tumor control achieved with IR + ACPP-AZD0156 was lost following CD8⁺ T-cell depletion, further confirming the reliance of AZD0156 radiosensitized tumor control on adaptive immunity, rather than direct cytotoxicity alone. These data are in alignment with emerging reports that DDR inhibitors can act as radio-immunosensitizers, particularly when combined with immune checkpoint inhibitors[36, 38, 39, 57, 62]. Finally, we demonstrated that ACPP-AZD0156 synergizes with PD1 blockade, resulting in durable tumor control in the otherwise immunotherapy-resistant B16 model. This is particularly encouraging given that many tumors exhibit primary resistance to immune-checkpoint blockade due to poor immune infiltration or exclusion[40, 63]. Thus, by promoting immunogenic cell death, leukocyte activation, and CD8⁺ T-cell recruitment, our spatially restricted platform may help convert immunologically “cold” tumors into a more immune-responsive state (**Figure 7**).

**Figure 7.**
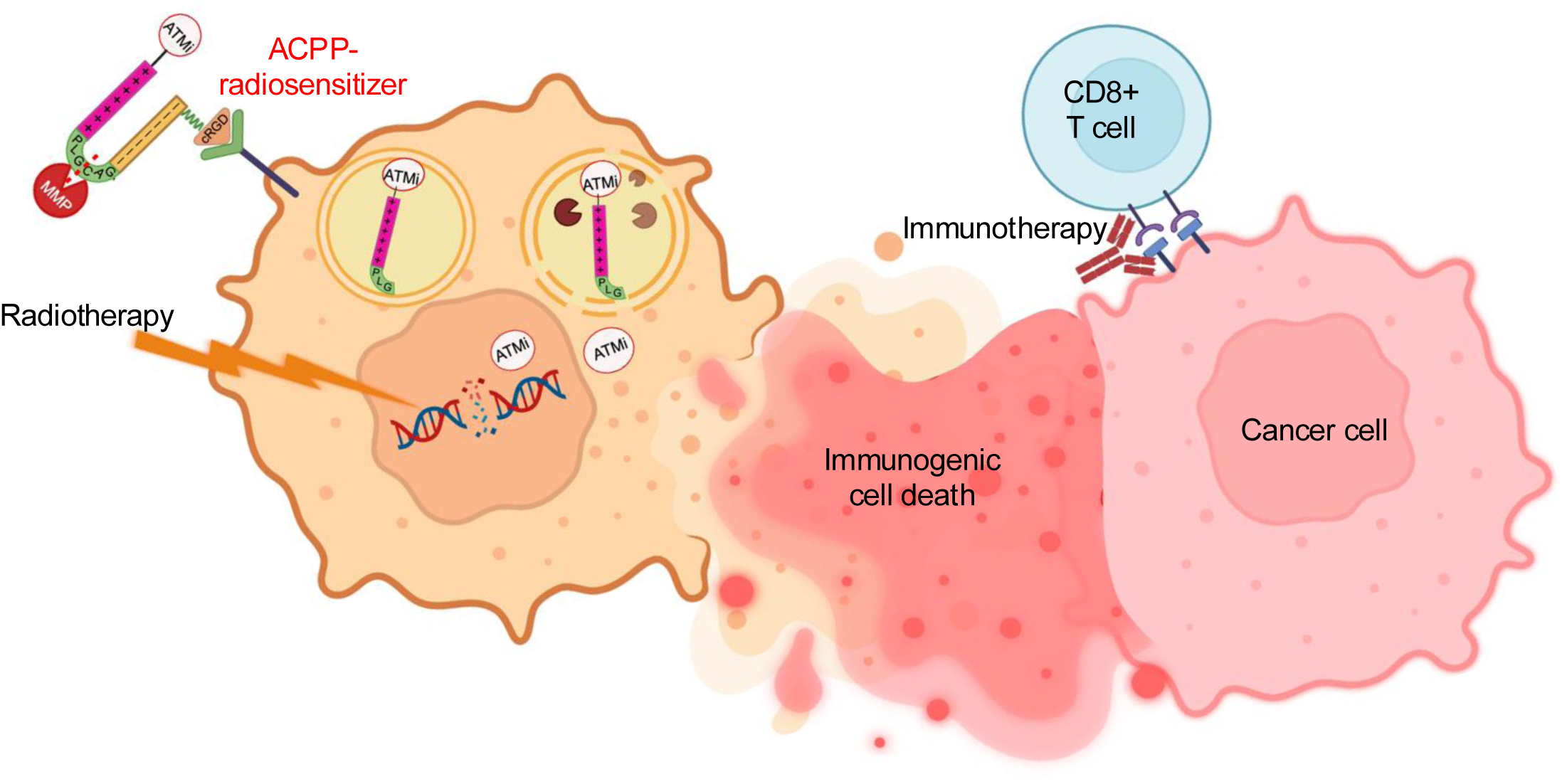
Activatable cell penetrating peptide targeted tumor radiosensitization to potentiate immunotherapies. Schematic model illustrating mechanism of targeted radiosensitization. Tumor-restricted activation of ACPP-AZD0156 by MMP2/9 results in spatially restricted ATM inhibition that enhances radiation-induced immunogenic cell death and CD8+ T-cell dependent tumor control and synergistic integration with immune checkpoint blockade.

In summary, our study provides preclinical proof-of-concept for tumor-targeted radio-immunosensitization strategy that integrates spatial ATM inhibition with immune modulation. Our ACPP scaffold permits the incorporation of diverse therapeutic payloads, including other DDR inhibitors such as ATR, DNA-PKcs, or PARP inhibitors, enabling precision radiosensitization integrated with immunotherapy depending on cancer-specific vulnerabilities. Together, our findings support further development of peptide-drug conjugate strategies as precision tools to enhance radiotherapy efficacy while promoting durable anti-tumor immune response.

## Supporting information

Differentially expressed genes summary

## Acknowledgements

This work was supported by the National Institutes of Health CA215081 (S.J.A.), CA268513 (S.J.A.), P30CA23100 (Microscopy Core, Moores Cancer Center), S10OD018499 (Flow Cytometry Core Facility, La Jolla Institute), T32 CA121938 (K.D.). Next Generation Sequencing (NGS) Core Facility of the Salk Institute was supported by P30CA014195, P30AG068635, the Chapman Foundation, and the Helmsley Charitable Trust. The authors thank Dr. Elsa Molina for technical assistance with NanoString studies. Schematic images in Figures 1A, 1B, 3A, 4A, 6E, and 7 were created using Bio.Render.com.

## Authors Contributions Statement

K.D., S.R.A., and S.J.A. conceived and designed the studies.

K.D., M.M.A., J.L., M.F.C., M.M.C., M.M., B.Z., and D.V.H. performed the experiments.

D.V.H. and S.R.A. synthesized peptide drug conjugates.

K.D., M.M.A., S.R.A., J.S.G. and S.J.A. interpreted the data.

S.J.A. supervised the studies.

## Data Availability

All data reported in this work are available upon reasonable request from the corresponding author.

## Supplementary Figure Legends

**Figure S1.**
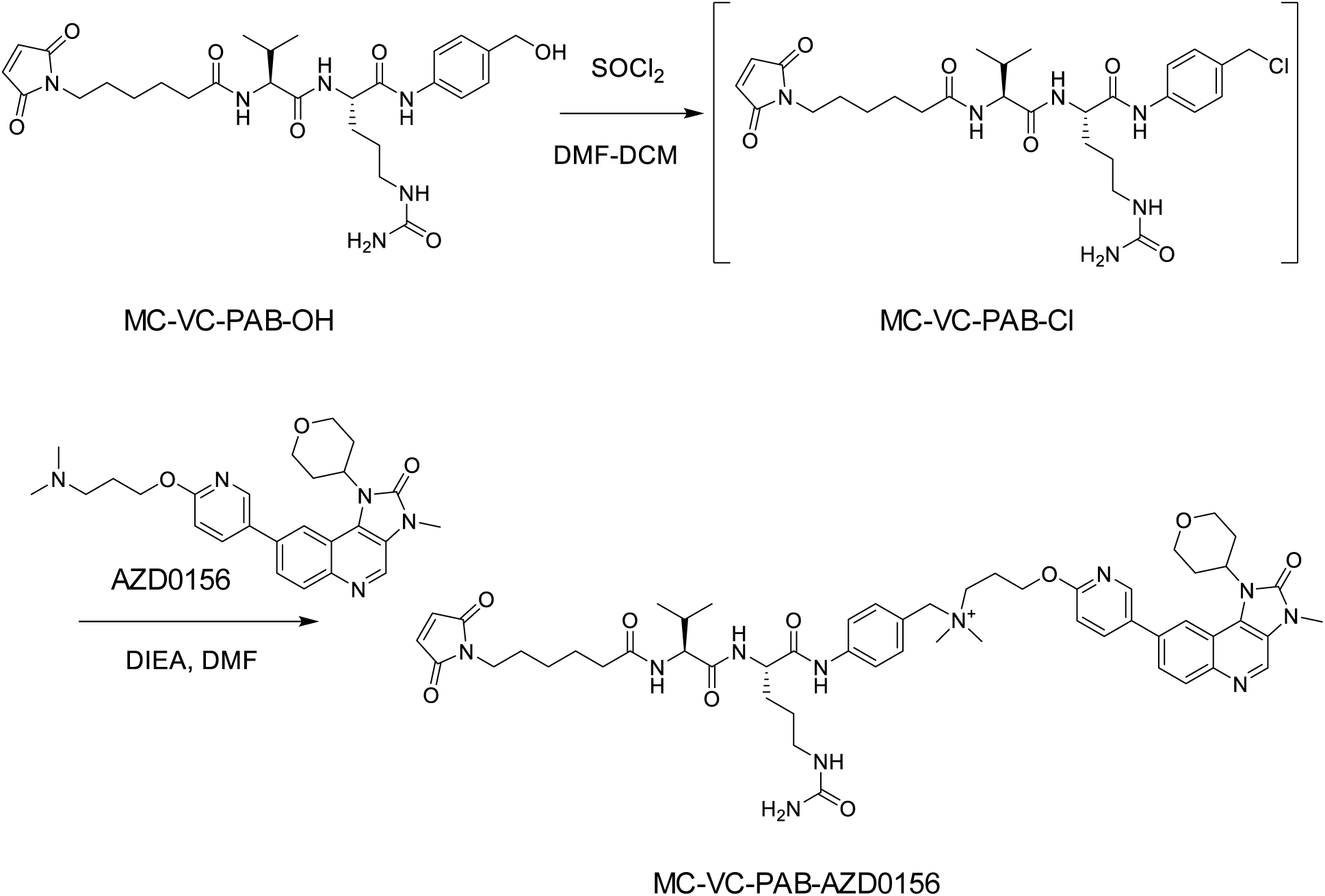
Synthesis of ATMi-linker conjugate. Synthesis of MC-VC-PAB-AZD0156 in which AZD0156 is attached to a maleimide cathepsin-cleavable self-immolative linker.

**Figure S2.**
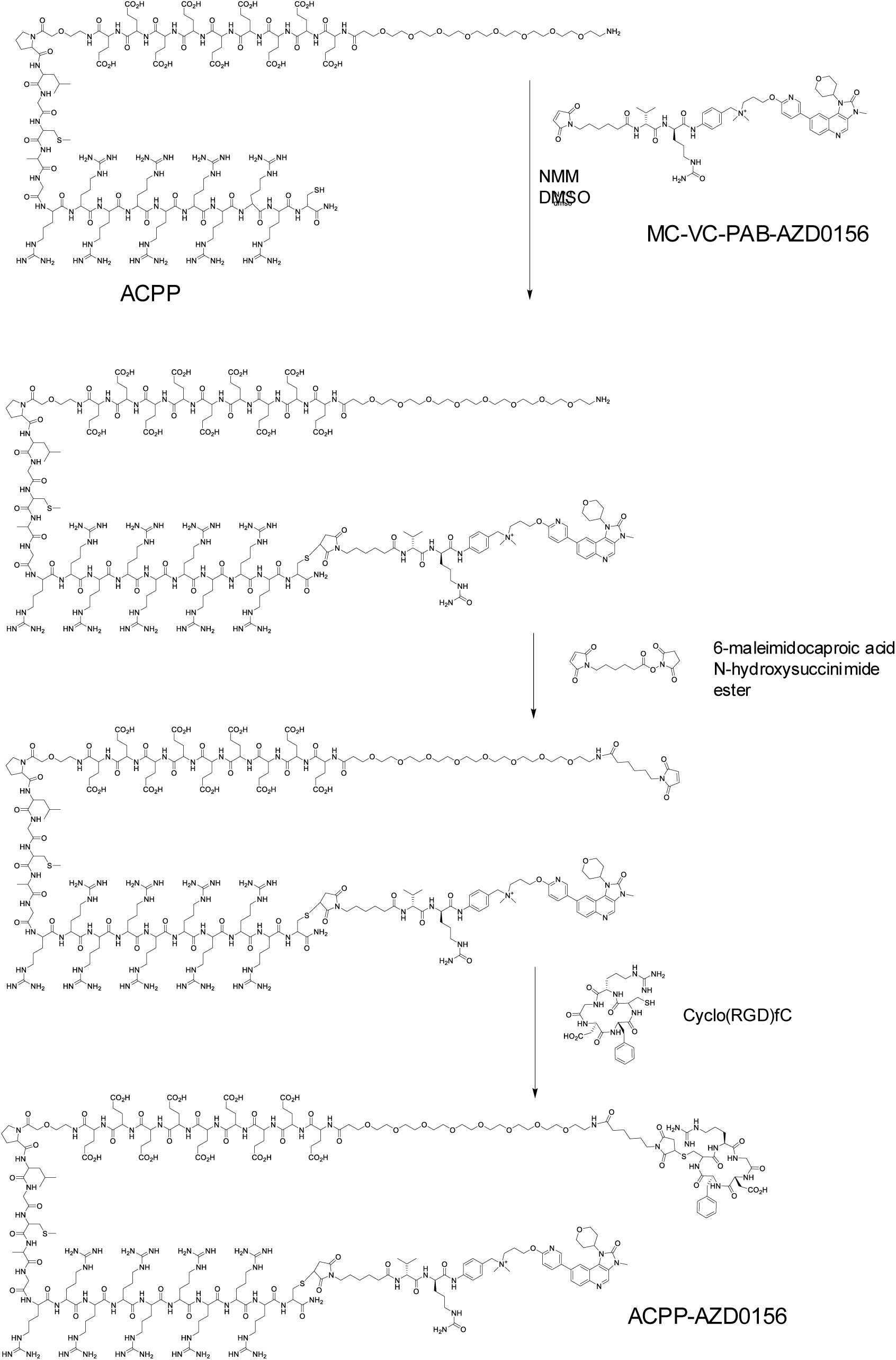
Synthesis of ACPP-ATMi conjugate. Synthetic scheme for activatable cell penetrating peptide conjugated to AZD0156. AZD0156 bound to MC-VC-PAB linker reacted with ACPP. cRGD reacted with ACPP conjugated AZ0156 to yield co-targeted cRGD-ACPP-AZD0156 (cyclo(RGD)fc-MC-HN-peg8-e_9_-oPLGC(Me)AG-r_9_-c-(MC-VC-PAB-AZD0156)-CONH_2_).

**Figure S3.**
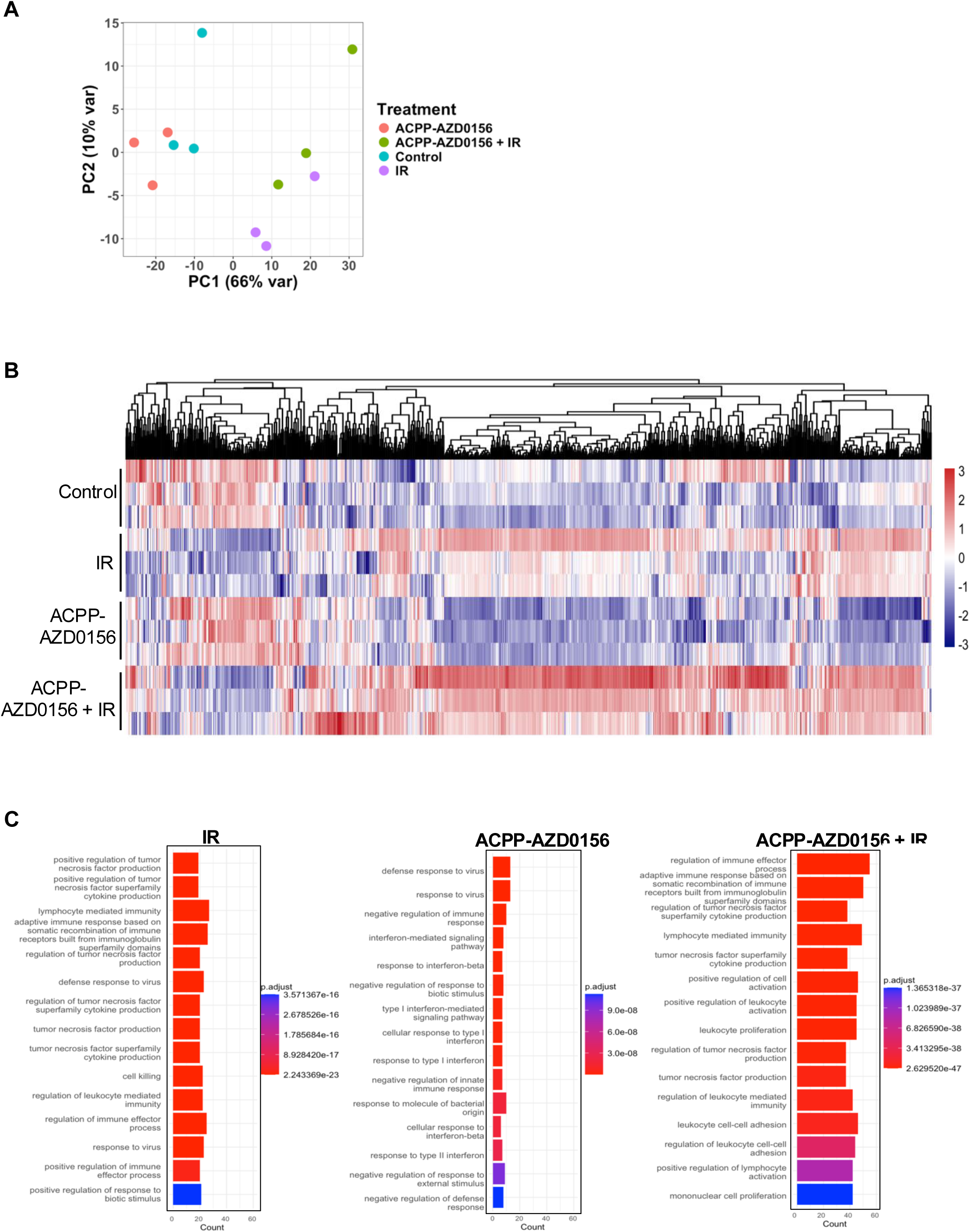
**A**) Principal component analysis plot based on normalized gene expression in B16 cells treated with 10 nM ACPP-AZD0156, ACPP-AZD0156 + IR or untreated (control) for 24 h, n = 3. RNA expression analyzed using NanoString PanCancer IO 360 Panel. **B**) Heatmap of gene expression for all samples as described in Figure S3a. The source data used in gene expression analysis is in the Source Data file. **C**) Pathway analysis results for IR (left), ACPP-AZD0156 (center), and ACPP-AZD0156 + IR (right) treated B16 cells vs control untreated samples, n=3. The top 15 biological processes from Gene Ontology that are enriched are displayed. Each bar depicts the enrichment scores (pvalues) and gene count within the corresponding gene set as bar height.

